# Differential effects of learned associations with words and pseudowords on event-related brain potentials

**DOI:** 10.1101/240945

**Authors:** Louisa Kulke, Mareike Bayer, Anna-Maria Grimm, Annekathrin Schacht

## Abstract

Associated stimulus valence affects neural responses at an early processing stage. However, in the field of written language processing, it is unclear whether semantics of a word or low-level visual features affect early neural processing advantages. The current study aimed to investigate the role of semantic content on reward and loss associations. Participants completed a learning session to associate either words (Experiment 1, N=24) or pseudowords (Experiment 2, N=24) with different monetary outcomes (gain-associated, neutral or loss-associated). Gain-associated stimuli were learned fastest. Behavioural and neural response changes based on the associated outcome were further investigated in separate test sessions. Responses were faster towards gain- and loss-associated than neutral stimuli if they were words, but not pseudowords. Early P1 effects of associated outcome occurred for both pseudowords and words. Specifically, loss-association resulted in increased P1 amplitudes to pseudowords, compared to decreased amplitudes to words. Although visual features are likely to explain P1 effects for pseudowords, the inversed effect for words suggests that semantic content affects associative learning, potentially leading to stronger associations.

**Highlights:**

- Neural mechanisms of gain/loss association to pseudowords and words were investigated
- Loss effects can be observed for the P1 component
- Words and pseudowords differ in the direction of loss effects
- Semantic content may play a role during word association
- Low-level visual features may play a role during pseudoword association

## 1. Introduction

Emotional valence leads to preferential processing of stimuli. These effects seem to occur already at an early level during neural processing, as recent research provided evidence for the occurrence of emotion effects in Event-Related Potentials (ERPs) already within the first 200 ms after stimulus onset, modulating the P1 component (Batty & Taylor, 2003; Bayer, Sommer, & Schacht, 2012b; Hofmann, Kuchinke, Tamm, Võ, & Jacobs, 2009; Rellecke, Palazova, Sommer, & Schacht, 2011; Scott, O'Donnell, Leuthold, & Sereno, 2009). These modulations occur both when a stimulus has an inherent emotional valence (e.g. Delplanque, Lavoie, Hot, Silvert, & Sequeira, 2004; Rellecke et al., 2011), as well as when the stimulus has been associated with a certain outcome due to reward or loss during conditioning. According to the dual competition model (Pessoa, 2009), emotional and motivational stimuli, similar to physically salient stimuli, are processed preferentially, leading to improved perceptual and behavioural responses towards them. Empirically, for example, gain-associated faces compared to zero outcome associated faces showed increases in P1 amplitude (Hammerschmidt, Sennhenn-Reulen, & Schacht, 2017) and boosted amplitudes of the Late Positive Complex (LPC) (Hammerschmidt, Kagan, Kulke, & Schacht, 2018; Hammerschmidt, Kulke, Broering, & Schacht, 2018). ERP modulations in the C1 time range were also reported for line gratings associated with threat-related pictures (Stolarova, Keil, & Moratti, 2005). Rossi et al. (2017) showed that meaningless symbols associated with monetary loss elicited larger C1 responses than neutral or reward-associated stimuli. Furthermore, in a study by Schacht, Adler, Chen, Guo, and Sommer (2012), formerly unfamiliar Chinese words were experimentally associated with monetary reward, loss, or no outcome. In a subsequent old/new decision task one day after the learning procedure, words associated with reward elicited enhanced P1 amplitudes. Together, these studies demonstrate that associative learning of complex unfamiliar visual stimuli influences sensory processing in the visual cortex. Neutral objects paired with emotional scenes elicited larger P1 and LPC (also termed Late Positive Potential, LPP, e.g. Schupp, Junghöfer, Weike, and Hamm (2004)) amplitudes compared to objects paired with neutral scenes (Ventura - Bort et al., 2016). Visual objects were also associated with positive taste, affecting LPC amplitudes (Blechert, Testa, Georgii, Klimesch, & Wilhelm, 2016). Neutral faces associated with negative smell elicited enhanced frontal and occipito-parietotemporal responses 50-80 msec and 130-190 msec after stimulus onset (Steinberg et al., 2012). Furthermore, faces that were associated with positive or negative auditory stimuli elicited early neural modulations as a function of valence, measured through MEG (Morel, Beaucousin, Perrin, & George, 2012). Additionally, faces associated with electric shocks induced changes in early prefrontal magnetoencephalography (MEG) activation (Rehbein et al., 2014) and in the LPC in ERPs (Rehbein et al., 2018) compared to non-associated faces. All these studies have in common that visual stimuli without any semantic meaning to the participant group were used, suggesting that modulations are likely related to the sensory encoding of a visual stimulus, leading to changes in early neural responses. In general, the P1 has been assumed to reflect sensory encoding of visual stimuli. It is modulated by the amount of attention allocated to a stimulus, showing an increase in amplitude for attended relative to unattended stimuli based on amplification of sensory information in the visual cortex (Hillyard & Anllo-Vento, 1998; Luck, Woodman, & Vogel, 2000).

However, in everyday life, not only images but also words are often associated with positive or negative emotions and early neural responses are modulated based on the inherent emotional valence of words (Bayer et al., 2012b; Rellecke et al., 2011). With regards to early neural responses, written words are an exceptional case, as most reading models assume that activations within the first 200 ms mainly reflect orthographic processing, while lexico-semantic features are accessed only later in the reading process (for a review, see Barber & Kutas, 2007). In contrast, there is a growing body of evidence suggesting that not only orthographic, but also lexico-semantic features can be accessed within the first 200 ms after stimulus onset (Hauk, Coutout, Holden, & Chen, 2012; Hauk, Davis, Ford, Pulvermüller, & Marslen-Wilson, 2006; Pulvermüller, Shtyrov, & Hauk, 2009; Rabovsky, Sommer, & Rahman, 2012). In particular, early modulations based on associated outcome have recently been observed for written pseudowords (Bayer, Grass, & Schacht, 2018). These findings are in accordance with evidence for fast activation of lexical information based on word frequency effects on fixation durations (Sereno & Rayner, 2003). The current set of experiments aimed at investigating whether words can be associated with gain or loss in an associative learning paradigm and to which extent semantic content effects these learning effects.

Taken together, previous findings provide two possible explanations for emotion effects in the P1 time range in response to words: First, they might be based on very fast semantic processing and thus corroborate the assumption that very fast recurrent feedback can modulate the activity of the visual cortex. Alternatively, associative learning of visual word features independent from the semantic system might contribute to P1 emotion effects. However, up to now, research does not allow for definite conclusions about the sources of early ERP effects driven by emotional content of written words. The neurocognitive mechanisms underlying these very early effects of emotional aspects from linguistic entities are in the main focus of the studies presented here.

Two further emotion-related ERP components will be investigated in the current study. Between approximately 200 to 350 ms after stimulus onset, emotional content causes the Early Posterior Negativity (EPN), a negative-going deflection in ERPs at posterior electrode sites (Bayer et al., 2012b; Kissler, Herbert, Peyk, & Junghofer, 2007; Palazova, Mantwill, Sommer, & Schacht, 2011; Schacht & Sommer, 2009a, 2009b). Concerning the localization of the EPN within the word recognition process, Schacht and Sommer (2009a) demonstrated that the EPN starts soon after lexical access has taken place, and therefore seems to reflect the activation of emotional *meaning* of written words (see also Palazova et al., 2011). At a later processing stage, starting around 400 ms after stimulus onset and lasting for several hundred milliseconds, emotional stimuli have been shown to increase the amplitudes of the LPC (e.g., Bayer, Sommer, & Schacht, 2010; Fischler & Bradley, 2006; Schacht & Sommer, 2009a). This centroparietal positivity has been suggested to reflect sustained elaborate processing of emotional stimuli (Cuthbert, Schupp, Bradley, Birbaumer, & Lang, 2000) and explicit decision processes (Schupp, Flaisch, Stockburger, & Junghöfer, 2006), presumably based on their intrinsic relevance for the observer (Schupp et al., 2004). This assumption is supported by a vast body of emotion-unrelated research on the P300 component – which the LPC is thought to be related to – suggesting that enhanced positive amplitudes in this time range are elicited by explicitly attended or task-relevant stimuli (Johnson, 1993).

In addition to emotion effects, associative learning has been demonstrated to induce Old/New effects (for an overview, see Rugg & Curran, 2007), which have also been observed in emotional association paradigms (Martínez-Galindo & Cansino, 2017), and can lead to larger P3 (also referred to as centro-parietal LPC), but not P1 amplitudes towards previously associated compared to novel stimuli (Bayer et al., 2018; Fritsch & Kuchinke, 2013). These effects will also be investigated in the current study to identify differences in the processing of acquired compared to novel stimuli.

Taken together, emotion effects in ERPs during visual word processing appear at the stage of perceptual processing (P1), sensory encoding of semantics (EPN), and higher-order stimulus evaluation (LPC). Especially in the case of P1 modulations, up to this point, there is no clear evidence concerning the possible sources of these effects, although two possible explanations – fast semantic access and associative learning of perceptual features – await scientific investigation. The present study aimed at investigating different origins of emotion effects in visual processing of linguistic stimuli and to specify their boundary conditions by associating words (Experiment 1) or pseudowords (Experiment 2) with gain, loss or no outcome. Both experiments only differ in whether the stimuli have an inherent (emotionally neutral) semantic meaning (Experiment 1) or not (Experiment 2), making it possible to directly investigate the effect that semantics have on associated outcome. Assuming that associated outcome affects early linguistic processing, gain- or loss-associated stimuli should elicit increased P1 amplitudes compared to neutral stimuli. Due to its relation to semantic processing, EPN effects were expected to be restricted to the processing of previously meaningful stimuli after associative learning (word condition) but should be absent in the case of entirely abstract although associated stimuli (pseudoword condition). Occipito-parietal LPC effects were expected to reflect conscious evaluation of acquired valence, independent of inherent semantics.

## 2. Methods

### 2.1 Participants

For Experiment 1 (words), data was collected in two sessions from twenty-five female students (*mean age* = 23.12 years; *SD* = 2.83 years, *range* = 19-31 years). One participant had to be excluded due to excessive artefacts within the EEG data, leading to a total of 24 subjects *(mean age* = 23.21 years; *SD* = 2.80 years, *range* = 19-31 years). For Experiment 2 (pseudowords), an independent group of 35 female students took part in two separate sessions *(mean age* = 24.71 years; *SD* = 4.13 years, *range* = 19-41 years). One subject had to be excluded due to prescription drugs affecting neural processes, 7 further subjects did not complete both sessions and 3 subjects were excluded due to excessive noise in the EEG data, leading to a total of 24 subjects *(mean age* = 23.96 years; *SD* = 3.09 years, *range* = 19-34 years). All participants were native German speakers recruited from the University of Göttingen, had normal or corrected-to-normal vision and reported no neurological or neuropsychological disorders or phobias. They were right-handed apart from four left-handers in Experiment 1 and two in Experiment 2.

Both experiments were conducted in accordance with the ethical principles formulated in the declaration of Helsinki and approved by the ethics committee of the University of Göttingen, Germany. Prior to both sessions, informed consent was obtained and participants were reimbursed with course credit or 8€ per hour plus performance-related bonuses (*M* = 6.93€ for words, *M* = 7.13€ for pseudowords, including a base pay of 3€).

### 2.2 Stimulus Material

A list of target stimuli used in both experiments is given in Appendix A. Stimuli for Experiment 1 consisted of 408 German nouns of neutral valence. They were selected from the Berlin Affective Word List Reloaded (Võ et al., 2009), based on the word features valence, arousal, imageability, word frequency and word length. Twenty-four words were selected as target words. For these words, the average of each word feature was within plus/minus one standard deviation of the mean, to ensure that the target words did not differ significantly from the 384 distractor words. The 24 target words were randomly split into six groups (for two blocks and three conditions) containing four words each with comparable word features. These six target groups were associated with one of three feedback conditions (gain, loss or zero outcome (neutral) feedback) in a counter-balanced way, so that every participant associated eight words with each feedback in the learning session. This counterbalancing ensures that the observed effects cannot be related to stimulus characteristics. The learning session consisted of two blocks separated by a pause. Within each block, four words out of every feedback condition (overall 12 words) were presented in sub-blocks that were repeated until the learning criterion was reached.

Target stimuli for Experiment 2 consisted of 24 disyllabic pseudowords following the phonological form consonant–vowel–consonant–vowel (e.g., foti, metu, bano). They were constructed in accordance with phonological rules of German and followed phoneme-grapheme correspondence. Stimuli were controlled with regard to sublexical bigram frequency (character bigram frequency, obtained from the dlex-database) and distribution of vowels. For the test session, 384 additional distractor pseudowords were constructed; analyses of all control variables listed above revealed no significant differences between target pseudowords and distractors, all *F*s(1,435) < 1.83, *p*s > .177.

### 2.3 Procedure

The procedure was kept identical in both experiments to allow for comparability. Both experiments consisted of two parts – a learning session and a test session. During the learning session, target stimuli were presented in random order within each block. One third of these stimuli were associated with monetary gain, monetary loss, or zero outcome (from here on referred to as neutral condition), respectively. In the learning session, participants were instructed to press one of three buttons – marked with the labels positive, negative, or neutral – with the index, middle, or ring finger of their dominant hand, within 5 s after stimulus onset. Participants were informed that some stimuli were coupled with monetary reward, some with loss and some did not have monetary consequences. In response to reward stimuli (gain-associated stimuli), pressing the correct button resulted in a gain of 20 cents, while only 10 cents were gained for erroneous responses. In response to loss-associated stimuli, 10 cents were lost when pressing the correct button while 20 cents were lost if an incorrect choice was made. Stimulus/outcome category-to-button assignment was counterbalanced between participants in all phases of the study. Missing responses resulted in a loss of 50 cents.

Trials started with a black fixation cross (font size 40pt. Arial) in the center of the light grey screen, visible for 500 ms, followed by one of 24 target stimuli, visible for a maximum of 5000 ms, which disappeared with the participant’s response. After a blank screen was visible for 1500 ms, a feedback stimulus was displayed for 1000 ms in the centre of the screen. Feedback stimuli consisted of grey circles with the amount of money gained or lost displayed inside, varying in font colour, signalling monetary gain (green), loss (red), neutral outcome (black), or timeout (blue), with a fixed font size of 45 pt. A fixed intertrial interval of 2000 ms followed the feedback, after which the next trial started. All 12 stimuli were presented once within each sub-block. After each sub-block, participants were informed about their current money balance. In order to successfully reach the learning criterion, participants needed to be correct in 48 out of the last 50 trials (moving window); after reaching this learning criterion, the learning session was terminated.

The test session took place one to two days after the learning session. It consisted of 16 sub-blocks. Participants had the opportunity to take short breaks after every sub-block with a longer break after eight sub-blocks. Each sub-block contained all 24 learned stimuli, presented in random order, with the same amount of unfamiliar, new distractor stimuli. In total, there were 384 distractors. Each trial started with a fixation cross visible for 500 ms followed by either a target or distractor word, visible for 1500 ms. Participants had to decide if the word was known from the learning session (target) or not (distractor) and press a button in response. No feedback was provided. The fixed inter-trial-interval was 2000 ms.

### 2.4 Data recording and pre-processing

The experiments were run using Presentation^®^ Software (Neurobehavioral Systems), which also recorded behavioural data. During the test session, the electroencephalogram (EEG) was continuously recorded from 64 active Ag-AgCl electrodes mounted in an elastic head cap (Easy Cap^™^) according to the extended international 10-20 system (Pivik et al., 1993). A BiosemiActiveTwo AD-Box amplified scalp voltage signals, which were digitized with 24 bits at a sampling rate of 512 Hz and amplified with a band pass filter of 0.16 – 100 Hz. Electrode offsets were kept at a threshold of +/-30 mV. Six external electrodes were applied to the left and right mastoids, the outer canthi (HEOG) and below both eyes (VEOG). Data was recorded with common mode sense (CMS) active electrode as reference and the driven right leg (DRL) passive electrode as ground electrode. Offline, the EEG was processed using Brain Vision Analyzer ^®^ Software. The continuous data was re-referenced to average reference. The HEOG and VEOG electrodes were used for correction of blink artefacts via Surrogate Multiple Source Eye Correction (MSEC; Ille, Berg, & Scherg, 2002), a built-in component of the BESA software (Brain Electrical Source Analysis, MEGIS Software GmbH). The data was downsampled to 500 Hz. A high-pass filter of 0.016 Hz, a low-pass filter of 40 Hz and a Notch filter of 50 Hz were applied. Minimal delays in the data were corrected by moving the triggers. The EEG was segmented into epochs of 1250 ms, starting 250 ms before stimulus onset, and referred to a 200 ms pre-stimulus baseline. Epochs were discarded as artefact-contaminated when any amplitude exceeded +/-200 μV or when any voltage step exceeded 100 μV between adjacent sampling points. Only epochs associated with correct responses were included in the analyses. Time windows for further data analysis were based on previous research and visual inspection of the grand average data. Based on previous literature, the mean P1 amplitude was extracted between 80 and 120 ms after stimulus onset in an occipital electrode cluster including electrodes O1, O2 and Oz; the mean EPN amplitude was extracted between 250 and 300ms after stimulus onset in an occipito-parietal electrode cluster including electrodes O1, O2, P9, P10, PO7 and PO8. The mean occipito-parietal LPC amplitude was quantified in a time window between 400 and 600 ms after stimulus onset in an occipito-parietal electrode cluster including Pz, POz, PO3 and PO4. In order to investigate differences between associated stimuli and novel stimuli, the centro-parietal LPC amplitude was extracted in an electrode cluster including centro-parietal electrodes CPz, P1, Pz, and P2 between 400 and 600ms after target onset, based on previous literature (Werheid, Schacht, & Sommer, 2007).

### 2.5 Data analysis

In the current study, the learning phase crucially differs from the test phase in that the number of trials until the learning criterion is met may differ between participants in the learning phase but trial number is constant in the test phase. Furthermore, error rates are expected to change across trials during learning. To account for these differences, different statistical approaches were chosen for the learning and test phase. To analyse behavioural data from the learning phase, posterior distributions for the probability (the coefficient *p* of a Bernoulli distribution) to attribute the outcome category correctly were analysed. The amount of evidence for underlying differences in this probability between the outcome categories was quantified. Based on a criterion of non-overlapping 99 % simultaneous credible bands, the time points during which conditions differed (given as a proportion of time until the learning criterion is met) were determined.

For analyses of behavioural data during the test phase, error rates were averaged within conditions and reported in percent of incorrect responses. Behavioural data was analysed using mixed linear models to investigate the effect of outcome (gain-associated, loss-associated or neutral) and stimulus type (pseudowords or words) as fixed factors and subject ID as random effect on reaction times and error rates. Differences between familiar and novel stimuli were investigated using a linear mixed model to predict reaction time and error rates from the fixed factors Old/New (familiar or novel stimulus) and stimulus type and subject ID as random effect.

For ERP analyses, mixed linear models were computed to investigate the effect of outcome (gain-associated, loss-associated or neutral) and stimulus type (pseudowords or words) as fixed factors and subject ID as random effect on ERP amplitudes of the P1, EPN and occipito-parietal LPC. To estimate effect sizes of significant effects within the mixed models, *R^2^m* was computed using the r-squaredGLMM function of the MuMIn package. For main effects within mixed models, effect sizes were estimated by dropping the interaction and other factor. Significant effects were followed up using dependent-sample t-tests. As directional hypotheses were stated for the comparison of the neutral with both other conditions, one-tailed tests were used (one directional prediction), while two-tailed tests were used to compare the positive and negative condition. Cohen’s d was computed as effect size for t-tests. The mixed model including subject ID as random effect was compared to an identical model excluding subject ID by comparing the difference in - 2 Restricted Log Likelihoods of the models with the *χ2*-distribution, to investigate the effect of individual differences. Differences between familiar and novel stimuli were investigated using a linear mixed model to predict ERP amplitude of the P1 and centro-parietal LPC from Old/New condition (whether the word was previously associated during the learning session or newly introduced in the test session) and stimulus type (word or pseudoword).

## 3. Results

### 3.1 Learning phase

#### Performance

Learning curves are displayed in Figure 1. In the **word condition**, gain-associated words were learned faster, differing from neutral (zero-outcome associated) words between 36.1% and 72.5% and between 41.0% and 99.9% in block 2 and from loss-associated words between 51.7% and 81.9% in block 1 and between 35.5% and 47.0% in block 2. There was no difference between loss-associated and neutral words in block 1 but a short advantage for loss-associated words between 81.5% and 99.9% in block 2.

**Figure 1.**
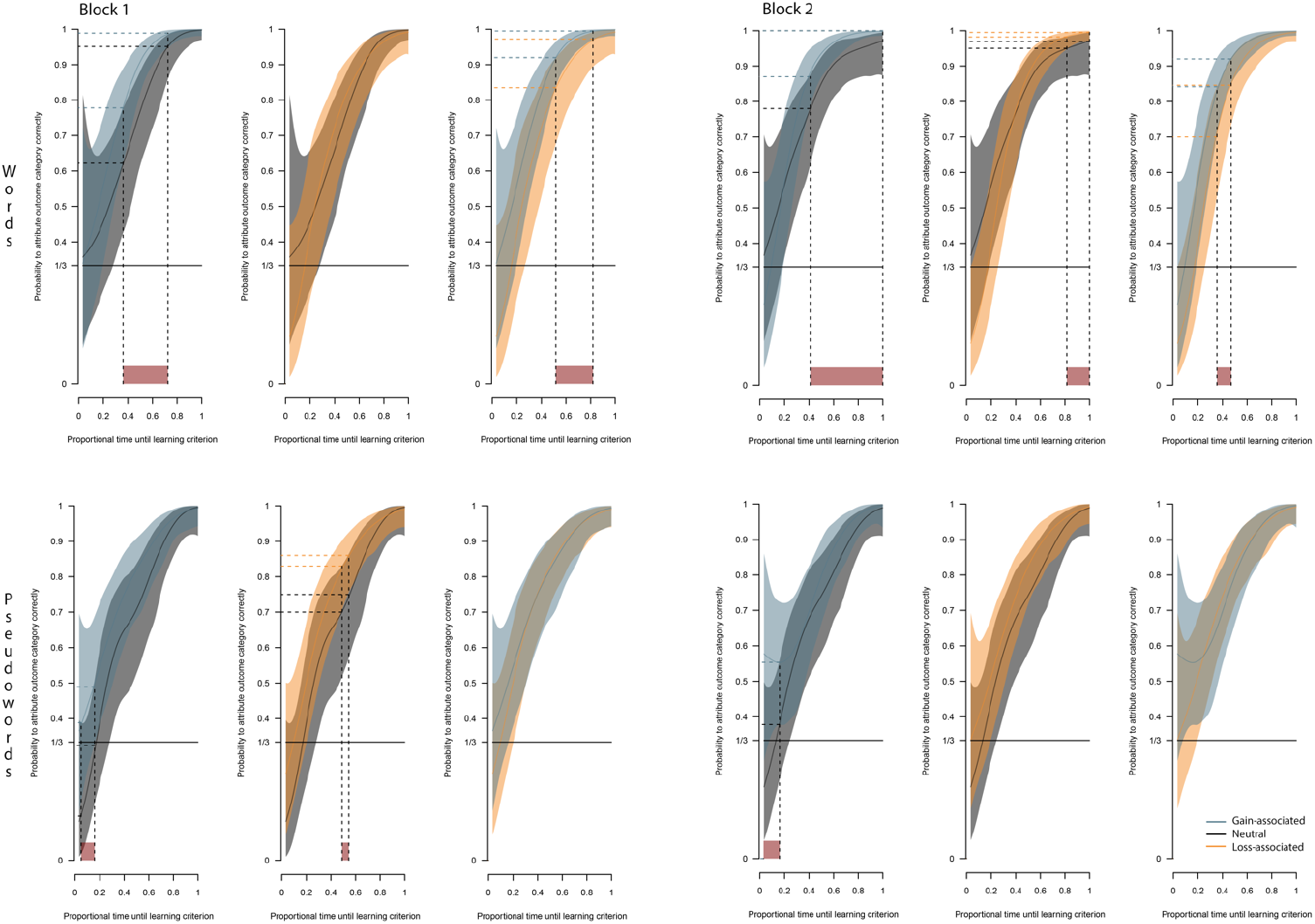
Posteriori mean probabilities to attribute the outcome category correctly as a function of proportional time until the learning criterion was met, separately for outcome categories (gain, loss, neutral). Differences between outcome categories outside the 99% confidence interval are illustrated by red areas and marked with horizontal dashed lines. The different panels show findings for block 1 (left) and block 2 (right) in the word (top) and pseudoword (bottom) condition.

In the **pseudoword condition**, gain-associated pseudowords were learned faster than neutral ones between 5% and 15.9% in block 1 and between 3% and 16.1% in block 2. Loss-associated pseudowords were learned faster than neutral pseudowords between 49.0% and 54.5% in block 1 but not in block 2. Gain-associated and loss-associated learning curves did not differ.

#### 3.1.2 Test phase

##### Performance

Figure 2 displays mean reaction times and error rates. Reaction times were significantly affected by outcome, *F*(2, 92) = 6.81, *p* = .002, *R^2^m* = 0.008, and stimulus type, *F*(1, 46) = 7.01, *p* = .011, *R^2^m* = 0.124, but there was no significant interaction, *F*(2, 92) = 2.57, *p* = .082. Percentages of errors showed no significant outcome, *F*(2, 92) = 2.50, *p* = .088, stimulus type, *F*(1, 46) = 2.56, *p* = .117, or interaction effect, *F*(2, 92) = 1.07, *p* = .348.

**Figure 2.**
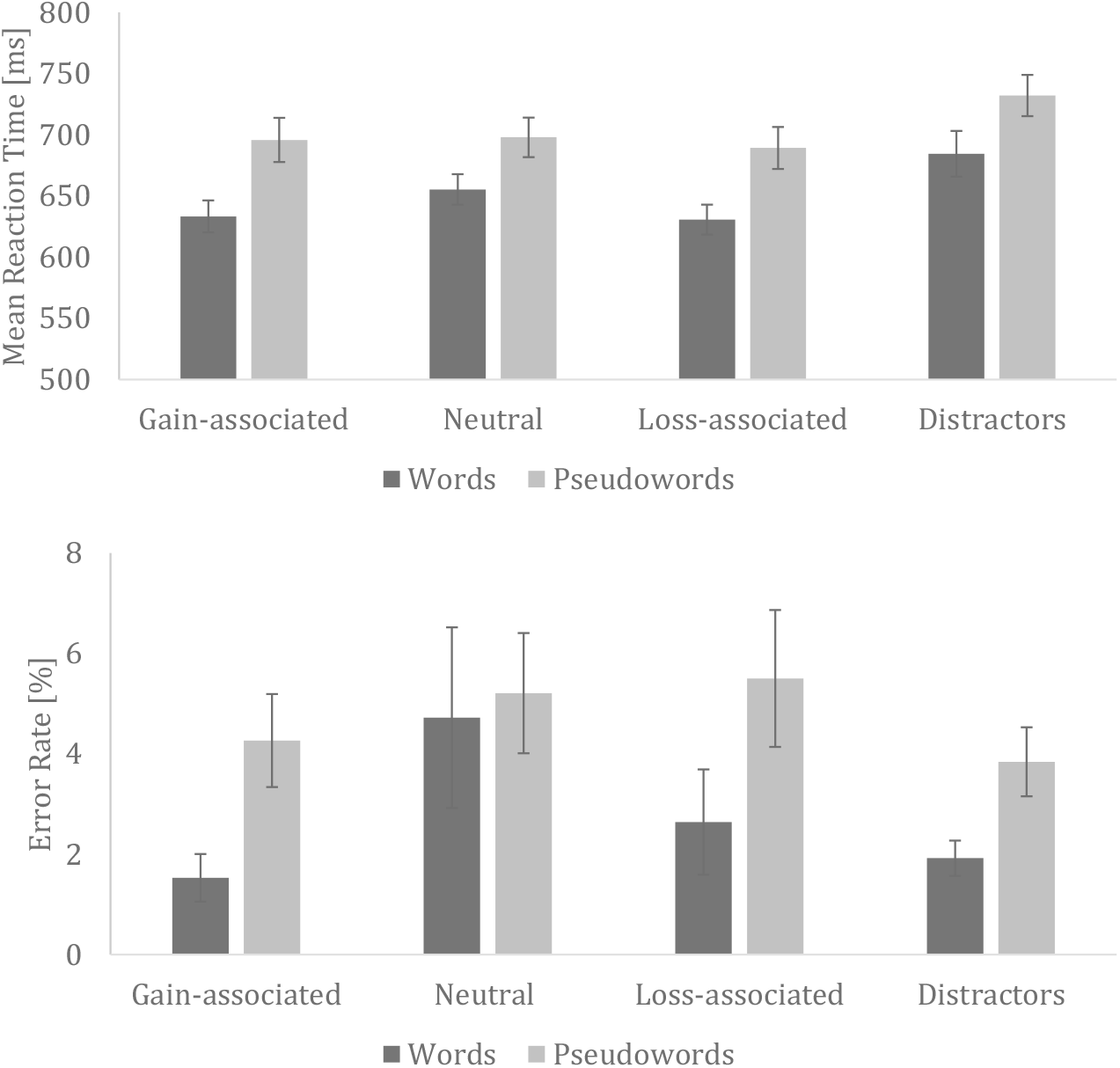
Mean reaction times (top) and error rates (bottom) to stimuli with associated outcome (gain-associated, neutral, loss-associated) and distractors in the test session. Error bars depict 2 *SE.*

Differences between familiar and novel stimuli were investigated using a linear mixed model to predict reaction time from Old/New and stimulus type. It showed a significant effect of Old/New, *F*(1, 46) = 30.35, *p* < .001, *R^2^m* = 0.060, and of stimulus type, *F*(1, 46) = 5.53, *p* = .023, *R^2^m* = 0.092, but no interaction, *F*(1, 46) = 0.21, *p* = .647. Percentages of errors showed a significant effect of stimulus type, *F*(1, 46) = 5.50, *p* = .023, *R^2^m* = 0.067, but no effect of Old/New, *F*(1, 46) = 2.54, *p* = .118, and no interaction, *F*(1, 46) = 0.004, *p* = .952. Hit rate (number of correct responses divided by trial number) showed a significant effect of stimulus type, *F*(1, 46) = 7.80, *p* = .008, *R^2^m* = 0.093, but no effect of Old/New, *F*(1, 46) = 2.81, *p* = .101, and no interaction, *F*(1, 46) = 0.00, *p* = 1.000. PR index (number of correct responses minus number of false positives) showed a significant effect of stimulus type, *F*(1, 46) = 5.25, *p* = .027, *R^2^m* = 0.101, but not of Old/New, *F*(1, 46) = 0.07, *p* = .796, and no interaction, *F*(1, 46) = 0.69, *p* = 0.412.

Separate analyses were conducted for each experiment to follow up on the effects described above.

##### Experiment 1: Words

Reaction times were significantly affected by associated outcome, *F*(2,46) = 10.00, *p* < .001, *R^2^m* = 0.033, due to faster reactions to loss-associated compared to neutral stimuli, *t*(23) = −4.24, *p* < .001, *d* = 0.865, and faster reactions to gain-associated than to neutral stimuli, *t*(23) = −3.23, *p* = .004, *d* = 0.660. Reaction times for loss-and gain-associated stimuli did not differ, *t*(23) = −0.48, *p* = .635.

There was a significant effect of Old/New, *F*(1, 23) = 14.07, *p* = .001, *R^2^m* = 0.082.

Mean error rates for all participants and conditions were at a low level of 2.80 %. Comparison of error rates between conditions revealed no main effect of outcome, *F*(2, 46) = 2.38, *p* = .104. There was no significant effect of Old/New, *F*(1, 23) = 1.27, *p* = .272. Hit rate, *F*(1, 23) = 1.56, *p* = .224, and PR index, *F*(1, 23) = 0.35, *p* = .559, showed no significant effect of Old/New.

##### Experiment 2: Pseudowords

Reaction times did not differ as a function of associated outcome, *F*(2, 46) = 0.814, *p* = .449. There was a significant effect of Old/New, *F*(1, 23) = 17.38, *p* < .001, *R^2^m* = 0.053.

Mean error rates for all participants and conditions were at an acceptable level of 4.42%. Comparisons of error rates between conditions revealed no main effect of outcome, *F*(2, 46) = 0.595, *p* = .556. There was no significant effect of Old/New, *F*(1, 23) = 1.27, *p* = .271. Hit rate, *F*(1, 23) = 1.27, *p* = .271, and PR index, *F*(1, 23) = 0.51, *p* = .484, also showed no significant effect of Old/New.

##### Neural findings — Effects of associated outcome

###### P1

There were no significant main effects of outcome, *F*(2, 92) = 0.37, *p* = .690, or stimulus type, *F*(1, 46) = 0.05, *p* = .825, but a significant interaction effect, *F*(2, 92) = 4.54, *p* = .013, *R^2^m* = 0.005, on P1 mean amplitudes. The model including subject ID as random effect (AIC = 492) had a significantly better fit than the identical model excluding subject ID (AIC = 711), *χ2* = 221, *p* < .001. Dependent sample t-tests comparing outcome conditions within each stimulus condition in the word condition showed a significant difference between the loss-associated and neutral stimuli, *t*(23) = - 3.89, *p* < .001, *d* = −1.59 (one-tailed, see Figure 3), but not between gain-associated and loss-associated stimuli, *t*(23) = 1.42, *p* = .169, *d* = 0.29 (two-tailed) or between the gain-associated and neutral stimuli, *t*(23) = −0.85, *p* = .203, *d* = −0.35 (one-tailed). In the pseudoword condition there was a significant difference between the loss-associated and neutral stimuli, *t*(23) = 1.85, *p* = .039, *d* = 0.76 (one-tailed), but no significant difference between the gain-associated and loss-associated stimuli, *t*(23) = −0.92, *p* = .370, *d* = −0.38 (two-tailed) or the gain-associated and the neutral stimuli, *t*(23) = 0.53, *p* = .302, *d* = 0.22 (one-tailed).

**Figure 3.**
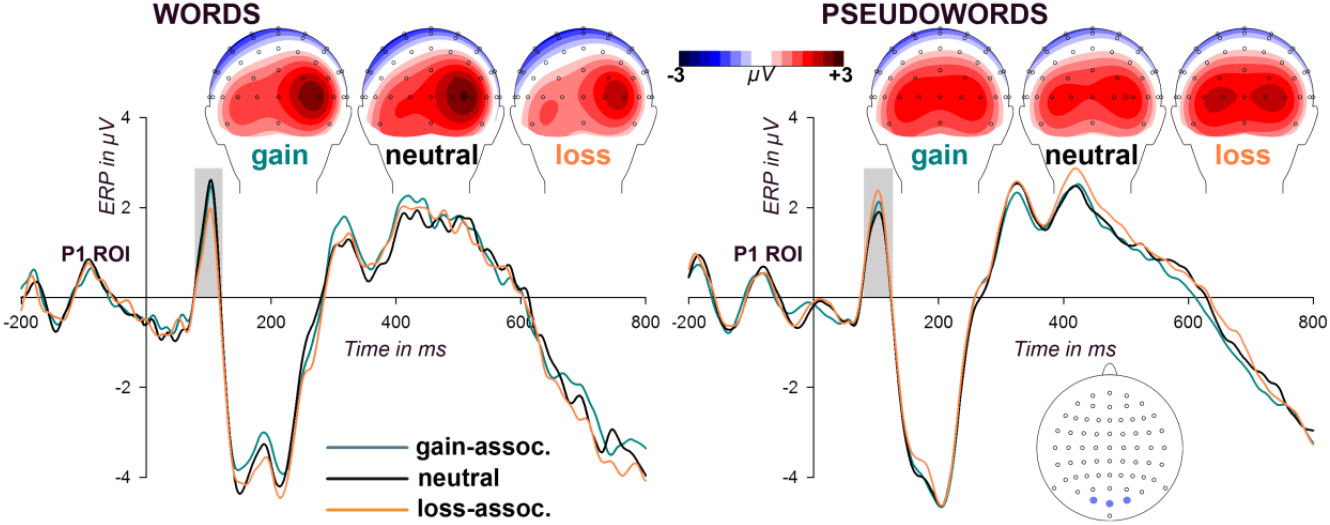
Effects of associated outcome on the P1 to words (left panel) and pseudowords (right panel). Grand average ERPs over P1 ROI electrodes, contrasted for gain-, loss-associated and neutral outcome, and time-locked to stimulus onsets. Inserted gray bars mark the time window of interest (80-120 ms), inserts highlight selected ROI electrodes (O1, O2 and Oz). Embedded heads depict the scalp distribution of ERP effects within the P1 time windows.

###### EPN

There were no significant main effects of stimulus type, *F*(1, 46) = 0.01, *p* = .906, or outcome, *F*(2, 92) = 1.32, *p* = .272, and no significant interaction, *F*(2, 92) = 2.00, *p* = .142, at the predetermined time window between 250 and 300ms after stimulus onset. The model including subject ID as random effect had a significantly better fit (AIC = 523) than the identical model excluding subject ID (AIC = 775), *χ2* = 254, *p* < .001.

Visual inspection of the wave forms suggested that, compared to the predetermined time-window based on previous literature, the EPN was shifted to an earlier time window between 150-250ms in the current study. A further analysis of mean amplitudes within this time-window showed that there were no significant main effects on early EPN amplitude of outcome, *F*(2, 92) = 1.11, *p* = .336, or stimulus type, *F*(1, 46) = 0.02, *p* = .883, and no interaction, *F*(2, 92) = 1.27, *p* = .287. The model including subject ID as random effect (AIC = 442) had a significantly better fit than the identical model excluding subject ID (AIC = 690), *χ2* = 250, *p* < .001.

###### Occipito-parietal LPC

There were no significant main effects of stimulus type, *F*(1, 46) = 0.19, *p* = .669, or outcome, *F*(2, 92) = 1.24, *p* = .294, and no significant interaction effect, *F*(2, 92) = 0.01, *p* = .988 on occipito-parietal LPC amplitudes. The model including subject ID as random effects (AIC = 528) was a significantly better fit than the identical model excluding subject ID (AIC = 787), *χ2* = 262, *p* < .001.

##### Distractor analyses

###### P1

A linear mixed model computing the effects of Old/New and stimulus type (word or pseudoword) on P1 amplitude showed no significant effect of Old/New, *F*(1, 46) = 0.41, p = .524, stimulus, *F*(1, 46) = 0.001, *p* = .980, or interaction, *F*(1, 46) = 3.49, *p* = .068.

###### Centro-parietal LPC

amplitudes showed a significant effect of Old/New condition, *F*(1, 46) = 141.80, *p* < .001, *R^2^m* = 0.197, but no effect of stimulus type, *F*(1, 46) = 1.35, *p* = .251, and no interaction, *F*(1, 46) = 0.57, *p* = .455. Follow-up t-tests showed that amplitudes were larger in the familiar than the distractor condition for words, *t*(23) = −8.32, *p* < .001, *d* = −3.40, and for pseudowords, *t*(23) = −8.60, *p* < .001, *d* = - 3.51 (Figure 4).

**Figure 4.**
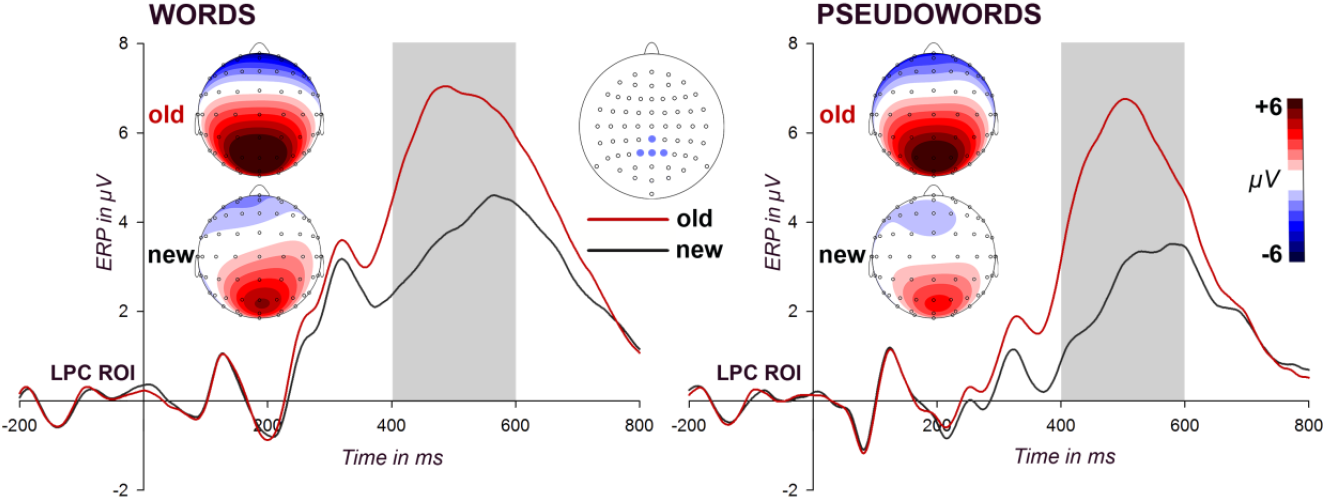
Grand average ERPs over centro-parietal LPC ROI electrodes, contrasted for learned (old) and new words (left panel) and pseudowords (right panel), and time-locked to stimulus onsets. Inserted gray bars mark the time window of interest (400-600 ms), inserts highlight selected ROI electrodes (CPz, Pz, P1 and P2). Embedded heads depict the scalp distribution of ERP effects within the centro-parietal LPC time windows.

## 4. Discussion

The current study investigated whether associated outcomes affect neural responses to word and pseudoword stimuli and what role semantic meaning plays in this process. On the behavioural level, gain-associated stimuli were acquired faster than neutral stimuli. In the test session, responses to stimuli associated with monetary gain or loss were significantly faster than to neutral stimuli in the word, but did not differ in the pseudoword condition. On the neural level, early responses showed significant effects of outcome with larger amplitudes to loss-associated compared to neutral stimuli in the pseudoword condition, but decreased amplitudes in the word condition. No outcome effects were detected on the EPN or occipito-parietal LPC. Finally, acquired stimuli led to faster reactions and increased centro-parietal LPC amplitudes compared to novel distractors.

### 4.1 Behavioural findings

The analyses of learning demonstrated that subjects acquired gain-associated words and pseudowords faster than neutral ones across all blocks. In line with previous research (Bayer et al., 2018; Hammerschmidt, Kulke, et al., 2018; Hammerschmidt et al., 2017; Rossi et al., 2017), this finding supports the assumption of a preferential processing of reward over loss on the behavioural level. Positive reinforcement (monetary gain) was used in the gain trials, but negative punishment (monetary loss) in the loss trials (Skinner, 1953). The loss, however, was decreased for correct compared to incorrect responses in the learning phase, simultaneously being a reward for correct learning. Gains and losses might therefore differ in their distribution to the observed effects. Advantages of gain over loss have previously been modelled based on behavioural data (Chapman, Gallivan, Wong, Wispinski, & Enns, 2015) and related to differences on the neural level (Hammerschmidt et al., 2017). In particular, the learning advantage for gain-associated stimuli in the current study started from a very early time during the learning process, possibly related to an early attentional draw towards gain. Once the pre-defined learning criterion was reached, the gain advantage levelled off. As participants’ reimbursement for the participation depended on task performance, the fact that a certain amount of monetary reward has already been gained might lower participants’ motivation towards gain over time, thus leading to the decreasing advantage for gain-associated stimuli. Since achieving the learning criterion required the successful learning of all associated outcome conditions, participants might have shifted their focus from the gain also to the other categories.

Behavioural findings from the old/new decision task in the test session confirmed that subjects successfully learned the stimuli during the learning session, as error rates were overall low. The lack of any modulation by associated outcome indicates similar familiarization of the stimuli across all outcome conditions. Furthermore, faster reaction times to associated stimuli than to novel distractors in both experiments confirm that the words and pseuodowords had been learned successfully. Effects of associated outcome on performance in the test session were restricted to reaction times to acquired words, with shorter reaction times to associated gain/ loss than neutral associations. One possible explanation is that words were more strongly associated with the respective outcome than pseudowords, as they convey a semantic meaning that can additionally prompt associations.

### 4.2 Neural findings

On the neural level, outcome effects on early evoked potentials appeared within the P1 time range, in line with previous findings (Bayer et al., 2018; Delplanque et al., 2004; Rellecke et al., 2011). This effect can only be related to the learned associations, but not to inherent stimulus properties, as the stimuli were assigned to each association group in a counterbalanced fashion, accounting for any innate valence of the respective stimuli. As the P1 is thought to reflect perceptual processing (Martínez et al., 2001), the current study ultimately demonstrates that early perceptual processing can be affected by associative learning. In line with the dual competition model (Pessoa, 2009, 2013), loss-association may change the motivational relevance of stimuli, leading to preferential processing in comparison to competing information.

The mechanisms underlying associated outcome effects are presumably complex and differed between words and pseudowords in the current study. An increase in P1 amplitude occurred for loss-associated compared to neutral pseudowords, while a decrease occurred for words. Increased P1 amplitudes have previously been reported for emotional pictures (Delplanque et al., 2004) and emotional facial expressions (Rellecke et al., 2011) and are thought to reflect increased attention to these stimuli (Heinze et al., 1994; Martínez et al., 2001). Pictures associated with an emotional context also induced larger P1 amplitudes (Ventura - Bort et al., 2016). Similar effects were found for pseudowords in the current study, with larger P1 amplitudes to loss-associated in contrast to neutral pseudowords. In the current study, the focus during stimulus construction was to create pseudowords matched for perceptual and psycholinguistic features. Accordingly, these stimuli should not be processed semantically, as a word stimulus, but rather as a visual object. In a similar study, Bayer et al. (2018) associated written and spoken pseudowords with positive (gain), negative (loss) or neutral outcome. They demonstrated that early ERP effects only occurred for visual stimuli that were also associated within the visual domain, but not for those associated in the auditory domain, highlighting the relevance of the visual shape during associative learning. Similarly, a study associating Chinese words with specific outcomes (Schacht et al., 2012) showed increases in early visual responses to stimuli with a gain-associated compared to neutral stimuli. In Western societies, neither Chinese characters, nor constructed pseudowords should have an inherent semantic meaning, leading to similar effects as for other non-semantic visual objects like faces and pictures. Furthermore, familiarity might play a role in early neural response modulation, as the overall combination of phonemes and graphemes may be more familiar for words than pseudowords. Therefore, current findings are in line with previous research showing an advantage of negatively associated stimuli if they were unknown (abstract) characters with a shape unfamiliar to the participants (Rossi et al., 2017). For words, however, we showed an opposite effect to previous studies using visual stimuli, suggesting that the semantic content of these words crucially affects early neural responses and their modulation through association. In fact, decreased P1 amplitudes have previously been reported for inherently negative compared to neutral words displayed in familiar font types (Kuchinke, Krause, Fritsch, & Briesemeister, 2014), suggesting that the effects due to associated outcome with words in the current study may be comparable to effects of inherent valence of words.

Kuchinke et al. (2014) showed that word features affect P1 modulations for familiar words. It is possible that pseudowords are even more strongly processed based on their visual features, while words are processed more holistically, including their semantic content. An early study by Johnston and McClelland (1974) demonstrated that single letters within a random letter string were better recognized if only the target letter was attended, while single letters within a word were better recognized if the full word was attended. Furthermore, the Stroop effect (Stroop, 1935) suggests that words are automatically processed holistically, interfering with their feature identification (for a review, see MacLeod, 1991). Low-level visual features were shown to have a greater effect on priming when pseudowords were used than when real words were used (Bowers, 1996), highlighting that visual features play a greater role for pseudowords than words. Taken together with the findings from the current study, unfamiliar pseudowords might elicit more feature-based processing, while familiar words might automatically be processed holistically, together with their semantic meaning, leading to similar findings for pseudowords (but not words) compared with findings from other unfamiliar stimuli.

Note that different groups of participants were recruited for both studies, though sampled from the same subject pool (student population of the same town). These participant groups may have differed in respect to their sensitivities to reward and loss. Although the current study cannot investigate effects of these differences, future research should measure participant traits, for example by using the BIS/BAS questionnaire (Carver & White, 1994).

Regarding the EPN component, we expected outcome effects to occur if the associated outcome would transfer to semantic meaning. This was not the case, as EPN amplitude did not differ as a function of associated outcome. Previous research showed EPN effects in tasks involving lexico-semantic processing (Bayer, Sommer, & Schacht, 2012a; Hinojosa, Méndez-Bértolo, & Pozo, 2010; Schacht & Sommer, 2009a) but not in tasks based on simple visual features (e.g. Rellecke et al., 2011) or Chinese words without semantic meanings to the subjects (Schacht et al., 2012). The lack of effects in the current study supports the idea that the EPN reflects the processing of word meaning (see also Palazova et al., 2011; Schacht & Sommer, 2009a) and that the stimuli in the current study were associated due to their visual features rather than their meaning. Another neural response of interest to emotional processing in the current study was the occipito-parietal LPC, an ERP component that has previously been linked to conscious evaluation of emotional valence as well as explicit emotion decision processes (Schacht & Sommer, 2009b; Schupp et al., 2006). Some previous associative learning paradigms showed the LPC to be affected by acquired stimulus valence (Hammerschmidt, Kagan, et al., 2018; Rehbein et al., 2018; Rossi et al., 2017; Schacht et al., 2012) while others reported no effects (Bayer et al., 2018; Hammerschmidt et al., 2017). The current study found no effects of associated outcome on occipito-parietal LPC amplitude, independent of semantic content, being in line with the latter. This mixed evidence leaves it open under which specific conditions (e.g., task requirements, stimulus types) modulations of late processing stages might occur.

An alternative explanation for the lack of effects of outcome category on later ERPs (EPN and LPC) is the potential extinction of associations during the test phase. During the learning phase, feedback related to the association category and response of the participant was presented in every trial. This unconditioned feedback stimulus was no longer presented in the test phase. Therefore, previous association effects may be attenuated during the test session. If this is the case, effects of outcome category might increase throughout the learning phase, but decrease throughout the test phase. This would lead to smaller effects when ERPs are averaged across the test phase than they may have been at the end of the learning session. Appendix B shows that behavioural responses did not significantly change between the first and the last block of the test session, suggesting that the effects remain fairly constant. Unfortunately, the current study only recorded ERPs during the test session, making it impossible to investigate whether stronger associations occured at the end of the learning phase. The development of effects over time should be investigated in future research.

The centro-parietal LPC responses previously related to associatively learned stimuli was significantly larger for learned compared to novel stimuli in the current study, being in line with previous research (Bayer et al., 2018; Fritsch & Kuchinke, 2013). This confirms that the stimuli have successfully been learned.

### 4.3 Divergence of behavioural and neural effects

Interestingly, the effects of outcome differed between learning behaviour and ERPs. Learning behaviour occurred faster for gain-associated stimuli, while ERPs were modulated by loss-associated stimuli. Both findings are in line with previous research (Bayer et al., 2018; Hammerschmidt et al., 2017; Rossi et al., 2017). The divergence indicates that different mechanisms play a role during the learning and the test phase. Participants seem to particularly respond to gain in the learning phase in order to optimise their reward, which can only be achieved through the correct behaviour. In contrast, the neural effects occurring in the test session are elicited by loss-associated stimuli, possibly related to their potential threat that needs to quickly and automatically be detected. Interestingly, MRI studies suggest that association of words with a negative context showed, amongst others, modulations in the lingual gyrus (Maratos, Dolan, Morris, Henson, & Rugg, 2001) and occipital regions (Smith, Henson, Dolan, & Rugg, 2004), which may be related to the posterior response measured in the current study. Positive context effects could only be detected in other brain regions (e.g. amygdala, hippocampus and dorsolateral prefrontal cortex, cf. Maratos et al., 2001; Smith et al., 2004), which should not be measurable in the P1 component. Follow up studies using MRI could reveal deeper brain regions involved in the observed modulations.

### 4.4 Conclusion

The current study aimed to investigate whether early response modulations are related to associations of visual features or fast semantic processing. On a behavioural level, associations with words had larger effects on reaction times than pseudowords. P1 enhancements to pseudowords without semantic content in the current study are likely to reflect associations with visual features. However, in an identical design, words involving a semantic meaning resulted in an opposite pattern regarding the P1 response. This indicates that semantic content, which is carried by words but not by pseudowords, might play a crucial role in associative learning. However, as no EPN modulations occurred, a later semantic processing of the words seems unlikely. It is unclear from the current study whether the effects are related to the familiarity of words compared to non-words, whether semantics directly influence early stage perception or whether semantic word meaning facilitates associations between visual features of a word and the associated outcome, leading to these P1 effects. However, words clearly differ from pseudowords regarding early neural mechanisms of outcome association in the current study and the underlying mechanisms should be further investigated in future research.

## Acknowledgements

This research was supported by the German Research Foundation (DFG; grant #SCHA1848/1-1 to AS). We would like to thank Holger Sennhenn-Reulen for providing scripts for the learning analyses and York Hagmayer for his statistical support.

## Declarations of interest

none

## Appendix A. Stimulus material

### Word stimuli

Gummi, Kohle, Sandale, Tonne, Bürste, Japan, Ritze, Stein, Abteil, Henne, Labor, Seil; Triangel, Halle, Kugel, Schwamm, Firma, Kasse, Rudel, Straße, Angel, Hocker, Person, Spule

### Pseudoword stimuli

bido, foti, gali, kego, bano, diru, fume, kowa, gipa, loni, sefu, weda; moke, niso, peli, wufa, bafe, gusa, melo, sogi, metu, nufe, rato, wola

## Appendix B. Block comparison

In order to investigate changes across blocks in the test phase, reaction time values were computed separately for the first and last block. Reaction time values for the first and last block are displayed Figure B.1. A mixed model predicting RT from Block and Old/New with Subject ID as random intercept showed a significant effect of Block, *F*(1, 141) = 202.52, *p* < .001, *R^2^m* = 0.216, of Old/New, *F*(1, 141) = 34.19, *p* < .001, *R^2^m* = 0.036, but no significant interaction, *F*(1, 141) = 0.000, *p* = .999.

A mixed model predicting RT from Block and Association with Subject ID as random intercept showed a significant effect of Block, *F*(1, 237) = 259.99, *p* < .001, *R^2^m* = 0.223, but no effect of association, *F*(1, 237) = 0.86, *p* = .356, and no significant interaction, *F*(1, 237) = 0.19, *p* = .664.

**Figure B.1.**
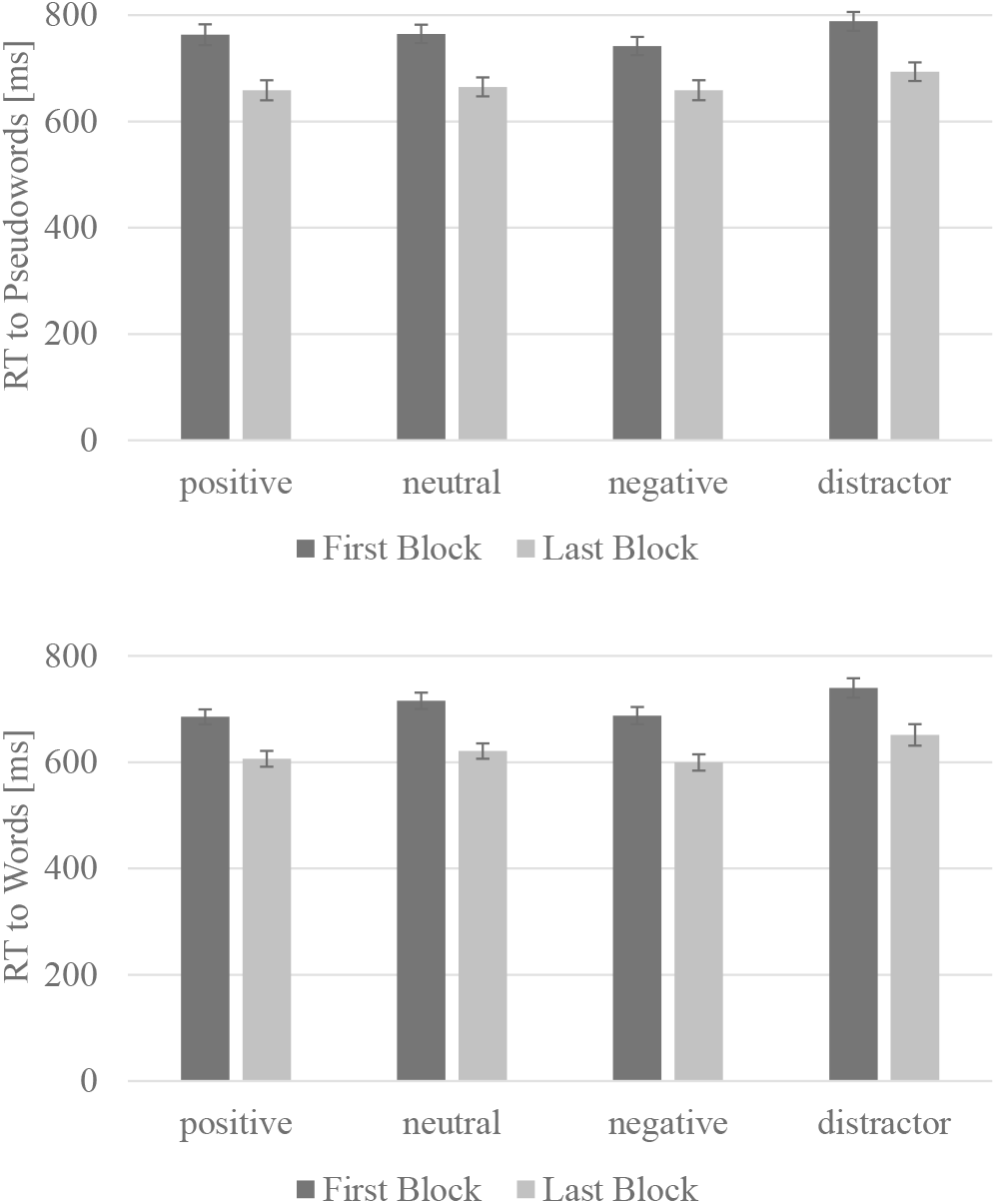
Reaction times (RT) to pseudowords (top) and words (bottom)

